# Endothelial Barrier Function is co-regulated at Vessel Bifurcations by Fluid Forces and Sphingosine-1-Phosphate

**DOI:** 10.1101/2020.08.18.256586

**Authors:** Ehsan Akbari, Griffin B. Spychalski, Miles M. Menyhert, Kaushik K. Rangharajan, Shaurya Prakash, Jonathan W. Song

## Abstract

Sphingosine-1-phosphate (S1P) is a blood-borne bioactive lipid mediator of endothelial barrier function. Prior studies have implicated mechanical stimulation due to intravascular laminar shear stress in co-regulating S1P signaling in endothelial cells (ECs). Yet, vascular networks *in vivo* consist of vessel bifurcations, and this geometry produces hemodynamic forces that are distinct from laminar shear stress. However, the role of these forces at vessel bifurcations in regulating S1P-dependent endothelial barrier function is not known. In this study, we implemented a microfluidic platform that recapitulates the flow dynamics of vessel bifurcations with *in situ* quantification of the permeability of microvessel analogues. Co-application of S1P with impinging bifurcated fluid flow, which was characterized by approximately zero shear stress and 38 dyn cm^-2^ stagnation pressure at the vessel bifurcation point, promotes vessel stabilization. Similarly, co-treatment of carrier-free S1P with 3 dyn cm^-2^ laminar shear stress is also protective of endothelial barrier function. Moreover, it is shown that vessel stabilization due to laminar shear stress, but not bifurcated fluid flow, is dependent on S1P receptor 1 or 2 signaling. Collectively, these findings demonstrate the endothelium-protective function of fluid forces at vessel bifurcations and their involvement in coordinating S1P-dependent regulation of vessel permeability.

## 1. Introduction

The endothelial cells (ECs) of small blood vessels, such as capillaries and post-capillary venules, form a semi-permeable barrier that control solute transport across the vessel wall.^[1]^ Maintenance of endothelial barrier integrity and permeability is crucial for regulating immune cell trafficking and tissue homeostasis.^[2]^ Accordingly, vascular barrier dysfunction underlies the pathogenesis of inflammation,^[3]^ atherosclerosis,^[4]^ cancer,^[5]^ and other diseases.^[6]^ Furthermore, heightened vessel permeability is a hallmark of pathological angiogenesis.^[7]^ Therefore, a fundamental understanding of the factors that help modulate endothelial barrier integrity is of great importance for restoring normal vascular function during disease conditions.

Sphingosine-1-phosphate (S1P) is a small and bioactive lysosphingolipid that signals through its family of G-protein coupled receptors.^[8]^ ECs are known to express three of the five known S1P receptors (S1PR1-3),^[9]^ and S1P exerts pleiotropic effects on EC proliferation, chemotaxis, and angiogenesis.^[10]^ S1P resides primarily in blood plasma at concentrations of 100-1000 nM^[11]^ and is an important regulator of endothelial barrier function.^[9, 12]^ Under normal physiological conditions, S1P associates primarily with two protein carriers or chaperones in order to be transported effectively through the bloodstream: high density lipoproteins (HDL; ∼65%) and albumin (∼35%).^[13]^ The effects of S1P on endothelial barrier function have been shown to be carrier-dependent. HDL-bound S1P, and to a lesser extent albumin-bound S1P, biases the activation of S1PR1,^[14]^ which is known to enhance endothelial barrier integrity.^[15]^ In contrast, activation of S1PR2 destabilizes endothelial junctions.^[16]^ Unlike carrier-bound S1P, carrier-free S1P engages all of the S1PRs with comparable affinity.^[17]^ Correspondingly, elevated levels of carrier-free S1P have been shown to induce a pro-inflammatory and atherogenic phenotype that is concomitant with compromised endothelial barrier function.^[18]^ Moreover, S1P production is upregulated in several human cancers compared with normal tissue,^[19]^ and antagonizing S1P signaling with targeted therapies^[20]^ have demonstrated anti-angiogenic effects in tumors.^[21]^

In addition to biomolecular signaling, fluid mechanical cues, such as laminar shear stress (LSS) that arises in straight regions of the vasculature, are known to be key regulators of vessel function.^[22]^ Interestingly, endothelial S1PR1 has been implicated as a mechanosensory molecule. LSS has been shown to upregulate endothelial S1PR1 expression levels^[23]^ and induce ligand-independent activation of S1PR1 that leads to suppression of sprouting angiogenesis and vessel stabilization *in vivo*.^[23c]^ Despite these findings, the role of hemodynamic forces in regulating the effects of free S1P on vessel function, especially in blood vessels that may be affected by fluid forces other than LSS, is poorly understood. Furthermore, while LSS predominantly arises in straight blood vessel segments, it is also important to consider that vascular networks are hierarchical branching structures that produce hemodynamic conditions at vessel bifurcations that are characteristically distinct from LSS. Specifically, at the base of vessel bifurcations, ECs are exposed to approximately zero shear stress but a finite pressure due to impinging flows that stagnate locally at the vessel bifurcation point.^[24]^

3-D microfluidic models of vascular function have been widely adopted for studying vascular biology and physiology under well-defined physical and chemical conditions *in vitro*.^[25]^ However, these models, including ones that studied the effects of S1P,^[10b, 26]^ feature either a single or two parallel linear channels lined with ECs.^[27]^ Consequently, to our knowledge the responses of S1P-signaling and flow dynamics at vessel bifurcations in modulating endothelial barrier function has not been investigated. In this study, we implemented our previously reported biomimetic microfluidic platform^[28]^ that uniquely combines *in vivo* settings (i.e., the flow dynamics at vessel bifurcations) with *in situ* quantification of the permeability coefficient hydraulic conductivity (*L*_*p*_) of microvessel analogues^[28a]^. We report that under static (i.e., no flow) conditions, treatment with carrier-free S1P (f-S1P) for 6 hours induces a ∼10-fold increase in *L*_*P*_ compared to untreated static control conditions. In comparison, co-treatment of f-S1P with impinging bifurcated fluid flow (BFF) (∼38 dyn cm^-2^ stagnation pressure and approximately zero shear stress) at the base of the bifurcation point (BP), decreases *L*_*P*_ significantly. Similarly, co-stimulation of f-S1P with physiological levels of LSS (∼3 dyn cm^-2^) in the branched vessel (BV) regions that are downstream of the BP also decreased *L*_*P*_ compared to f-S1P under static conditions. Furthermore, using pharmacological antagonists for S1PR1 (W146) and S1PR2 (JTE013), we found that flow-mediated stabilization of LSS, but not BFF, is mediated by these two S1P receptors. The findings reported here demonstrate the importance of fluid mechanical cues at vessel bifurcations in coordinating S1P-mediated endothelial barrier function.

## 2. Results

### 2.1. Microfluidic model of a bifurcating vessel for the study of S1P-dependent endothelial permeability

A biomimetic microfluidic model (**Figure 1**) was fabricated using poly(dimethylsiloxane) (PDMS) soft lithography, as previously reported^[28]^. Briefly, this microfluidic model consists of an inlet microchannel that bifurcates around a central extracellular matrix (ECM) compartment separated by PDMS microposts to form two smaller, equally wide microchannels (**Figure 1A**). The microfluidic model allows for the simultaneous application of BFF at the base of the BP, where the flow stagnates, and LSS in the branched vessels BV (**Figure 1Bi**). Furthermore, the microfluidic device allows for direct contact between human umbilical vein endothelial cells (HUVECs) and abluminal three-dimensional (3-D) collagen matrix at both the BP and the BV locations to examine the effect of the corresponding fluid forces on *L*_*P*_ (**Figure 1Bii, iii**). The microchannels were uniformly lined with a monolayer of HUVECs (**Figure 1Ci**), allowing for complete coverage of the ECM apertures at both BP and BV (**Figure 1Cii, iii**). Moreover, the microfluidic platform enables controlled administration of S1P in the microchannels under both static and flow conditions.

**Figure 1.**
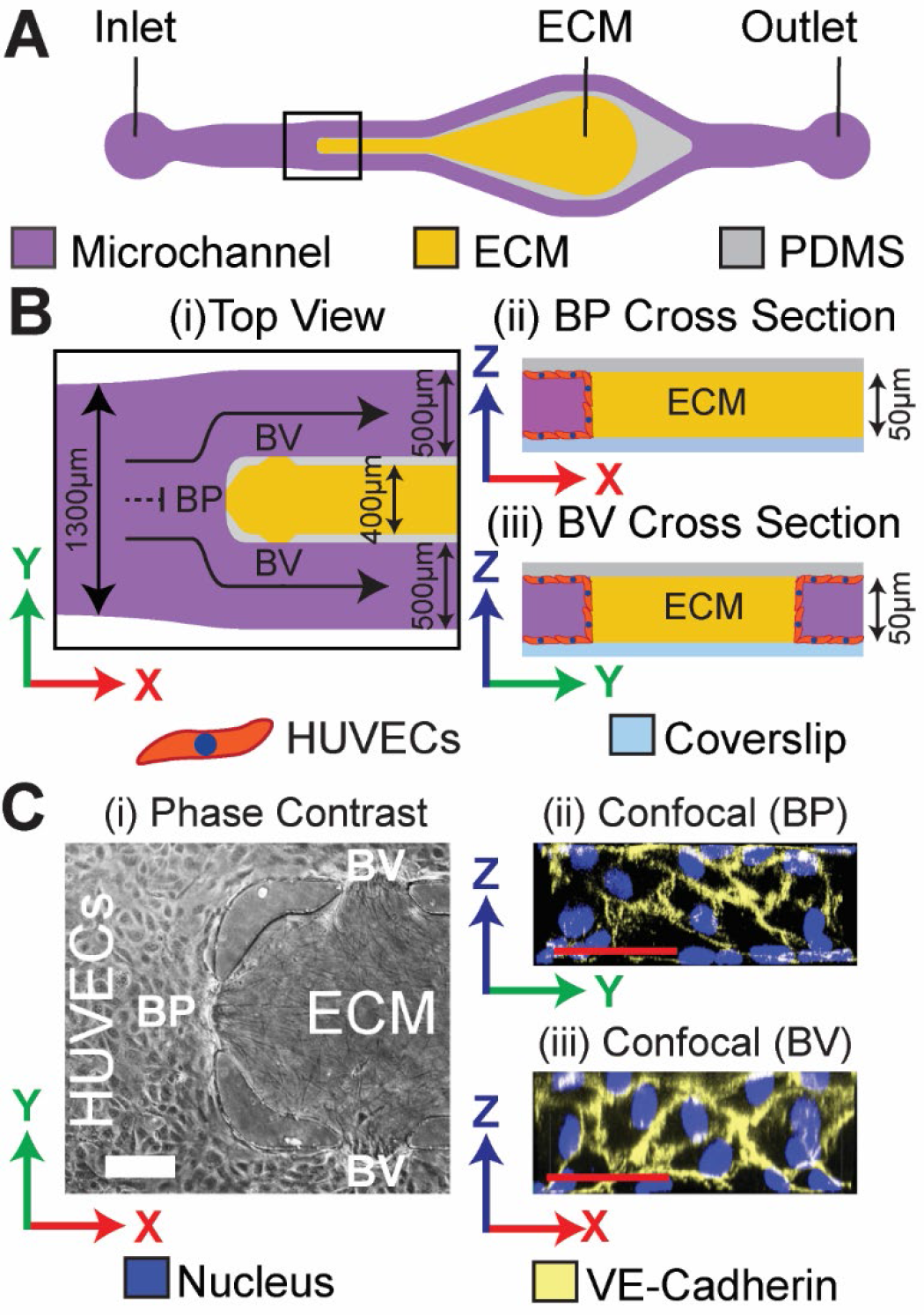
Biomimetic microfluidic model of vessel bifurcation for studying S1P-dependent *L*_*P*_. (A) The device top view schematic depicting the inlet channel bifurcating into two smaller channels around a central extracellular matrix (ECM) compartment. (B) The zoomed-in view of the bifurcation region (denoted by the black box in A). (i) Top view schematic depicts the laminar inflow stagnating on the base of the bifurcation point (BP) that results in application of bifurcated fluid flow (BFF, black dash line). Downstream of the BP, flow develops into two regions that are under laminar shear stress (LSS, black solid lines) in the branched vessels (BV). Moreover, the apertures included in the PDMS barrier that separates the central ECM compartment from the endothelial channels allow for the formation of the endothelial monolayer at: (ii) the BP, and (iii) in each BV on the ECM. (C) Representative (i) phase contrast and (ii) confocal immunofluorescence images of the BP fully seeded with a confluent monolayer of HUVECs that have formed well-defined adherent junction structures. White scale bar is 100µm. Red scale bars are 50µm.

### 2.2. Fluid mechanical forces associated with BFF and LSS attenuate endothelial permeability induced by carrier-free S1P

We first examined the effect of carrier-free S1P (f-S1P) on *L*_*P*_ under static or no-flow conditions in our microfluidic model. Treatment with 50 nM f-S1P for 6 hours resulted in a 3.6-fold increase in *L*_*P*_ (5.04×10^−4^ ± 1.00×10^−4^ cm·s^-1^·cmH_2_O^-1^, *p*<0.01) compared to the untreated and static condition (1.39×10^−4^ ± 0.21×10^−4^ cm·s^-1^·cmH_2_O^-1^), henceforth referred to as baseline *L*_*P*_ (**Figure S1**). Moreover, treatment with 500 nM f-S1P for 6 hours induced 10-fold increase in *L*_*P*_ (13.88×10^−4^ ± 2.54×10^−4^ cm·s^-1^·cmH_2_O^-1^, *p*<0.001) compared to baseline *L*_*P*_ (**Figures S1, 2B, 2D**). These results demonstrate a clear dose-dependent increase in *L*_*P*_ due to treatment of HUVECs with f-S1P. The induction of increased permeability of HUVECs with f-S1P at both of the tested concentrations (50 nM and 500 nM) is supported by previous *in vitro* reports of the pro-angiogenic effects of f-S1P.^[10a, 10b]^

Next, we examined the effects of co-stimulation of f-S1P with fluid flow on *L*_*P*_ at both the BP and BV apertures. The media perfusion flow rate was adjusted to produce 3 dyn cm^-2^ LSS in the BV regions of the microfluidic model, which is within the physiological range of shear stress in post-capillary venules (**Figure 2A**).^[29]^ This flow rate of 10 µL/min generates a 38 dyn cm^-2^ stagnation pressure with approximately average zero shear stress (**Figure 2A**) as reported previously.^[28a]^ Compared to the 500 nM f-S1P treatment under static conditions, co-application of 500 nM f-S1P with 38 dyn cm^-2^ SP and 3 dyn cm^-2^ LSS for 6 hours resulted in a significant decrease in *L*_*P*_ at the BP (0.78×10^−4^ ± 0.24×10^−4^ cm·s^-1^·cmH_2_O^-1^, *p*<0.01) and BV regions (0.87×10^−4^ ± 0.16×10^−4^ cm·s^-1^·cmH_2_O^-1^, *p*<0.001) respectively (**Figure 2B, D**). Our findings suggest that the fluid mechanical forces associated with 38 dyn cm^-2^ BFF and 3 dyn cm^-2^ LSS are endothelium-protective and counteract the induction of permeability due to f-S1P.

**Figure 2.**
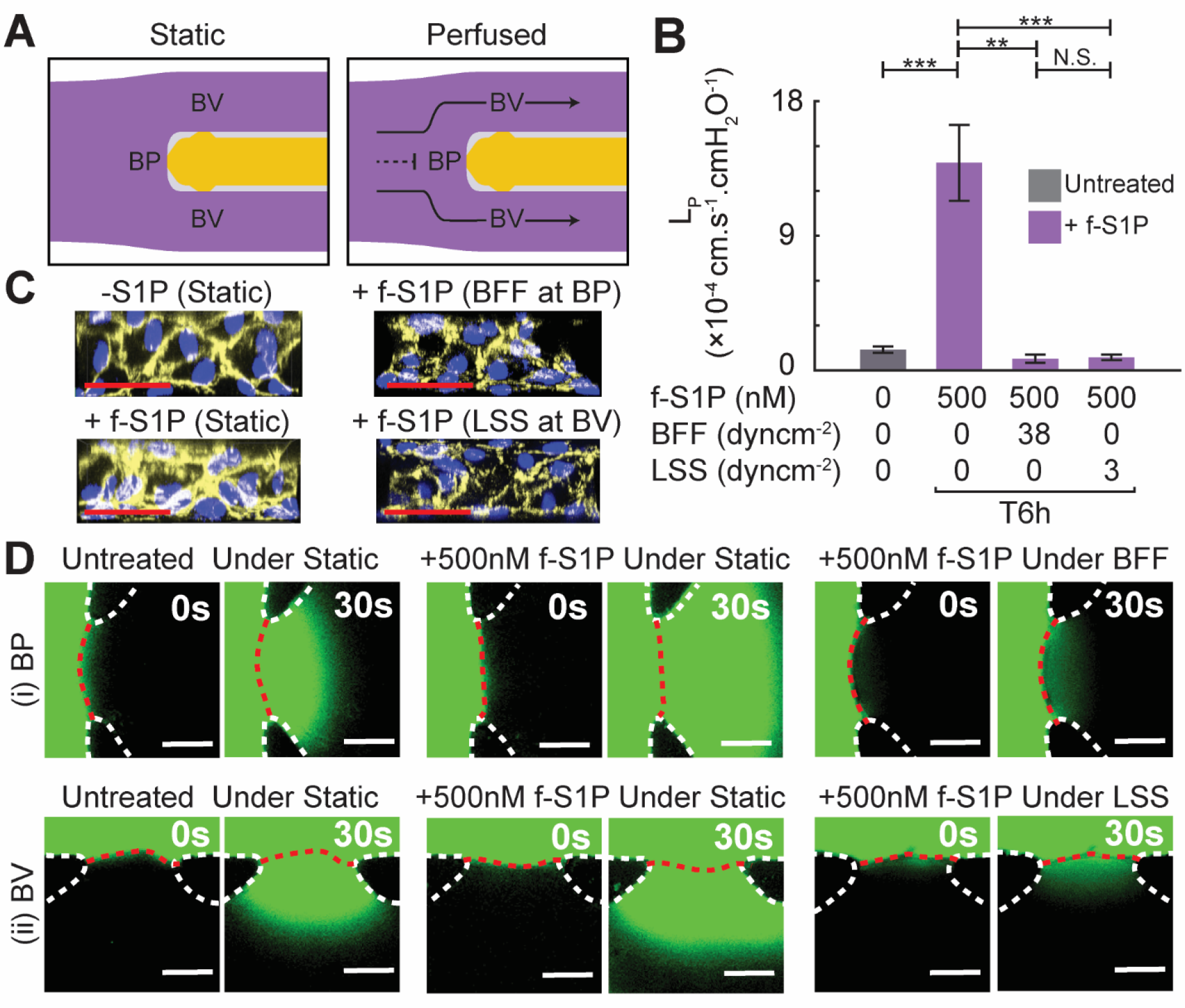
Application of BFF and LSS results in significant attenuation of increase in *L*_*P*_ induced by f-S1P. (A) Schematic of the experimental conditions to test the effect of BFF (black dash line) and LSS (black solid line) on S1P-dependent *L*_*P*_ compared to static control. (B) Quantitative response of HUVEC *L*_*P*_ to treatment with S1P under static condition compared to S1P co-applied with BFF or LSS. (C) Representative confocal images of VE-cadherin expression under each experimental test condition. Blue: HUVEC Nuclei, Yellow: VE-cadherin. (D) Representative epi-fluorescence images of FITC-Dextran extravasation rate to measure *L*_*P*_ at BP and BV after treatment under each experimental condition. Scale bars are 50µm. **: p < 0.01, ***: p < 0.001.

An important regulator of vascular permeability is the inter-endothelial junction protein VE-cadherin.^[30]^ Moreover, a previous study demonstrated the role of S1P in regulating the expression of VE-cadherin.^[15a]^ Therefore, we assessed VE-cadherin spatial expression patterns using immunofluorescence and *en face* confocal microscopy images of the endothelial monolayers at the BP and BV apertures under different f-S1P and perfusion conditions (**Figure 2C**). By visual inspection, treatment with f-S1P under static condition increased VE-cadherin expression levels at inter-endothelial junctions compared to the untreated static condition. In contrast, VE-cadherin expression at inter-endothelial junctions was similar or slightly reduced when f-S1P was co-applied with BFF and LSS (**Figure 2C**) when compared against the untreated static condition. These results suggest that the enhancement in endothelial barrier function due to BFF and LSS in the presence of f-S1P (**Figure 2B**) was independent of VE-cadherin localization to inter-endothelial junctions (**Figure 2C**).

### 2.3. Flow-mediated stabilization due to LSS, but not BFF, is dependent on S1PR1 or S1PR2 signaling

Previous reports suggest that the effect of S1P on endothelial barrier function is dependent on the relative activation of S1PR1 and S1PR2.^[8, 16, 31]^ These previous findings prompted us to study the involvement of S1PR1 and S1PR2 in regulating BFF and LSS mediated vessel stabilization (**Figure 3A**). We blocked S1P signaling with two widely used pharmacological antagonists with specificity for S1PR1 (W146) and S1PR2 (JTE013).^[12b, 16, 32]^ Under static conditions, pre-treatment with W146 followed by treatment with 500 nM f-S1P for 6 hours resulted in a significant decrease in HUVEC *L*_*P*_ (0.65×10^−4^ ± 0.21×10^−4^ cm·s^-1^·cmH_2_O^-1^, *p*<0.0001) compared to the f-S1P treated condition with no receptor inhibition (**Figure 3Ai,Bi**). This result suggests that activation of S1PR1 by f-S1P is involved with the induction of vessel permeability under static conditions. This observation was surprising as multiple studies have highlighted the role of S1PR1 signaling in stabilizing the endothelium.^[13b, 33]^ Under static condition, pretreatment with JTE013 followed by treatment with 500nM f-S1P also resulted in a significant decrease in *L*_*P*_ (0.41×10^−4^ ± 0.08×10^−4^ cm·s^-1^·cmH_2_O^-1^, *p*<0.0001) compared to the f-S1P treated condition with no inhibitor applied (**Figure 3Ai,Bi**). This result is in accordance with previous reports on the involvement of S1PR2 in S1P-induced endothelial barrier destabilization.^[16]^

**Figure 3.**
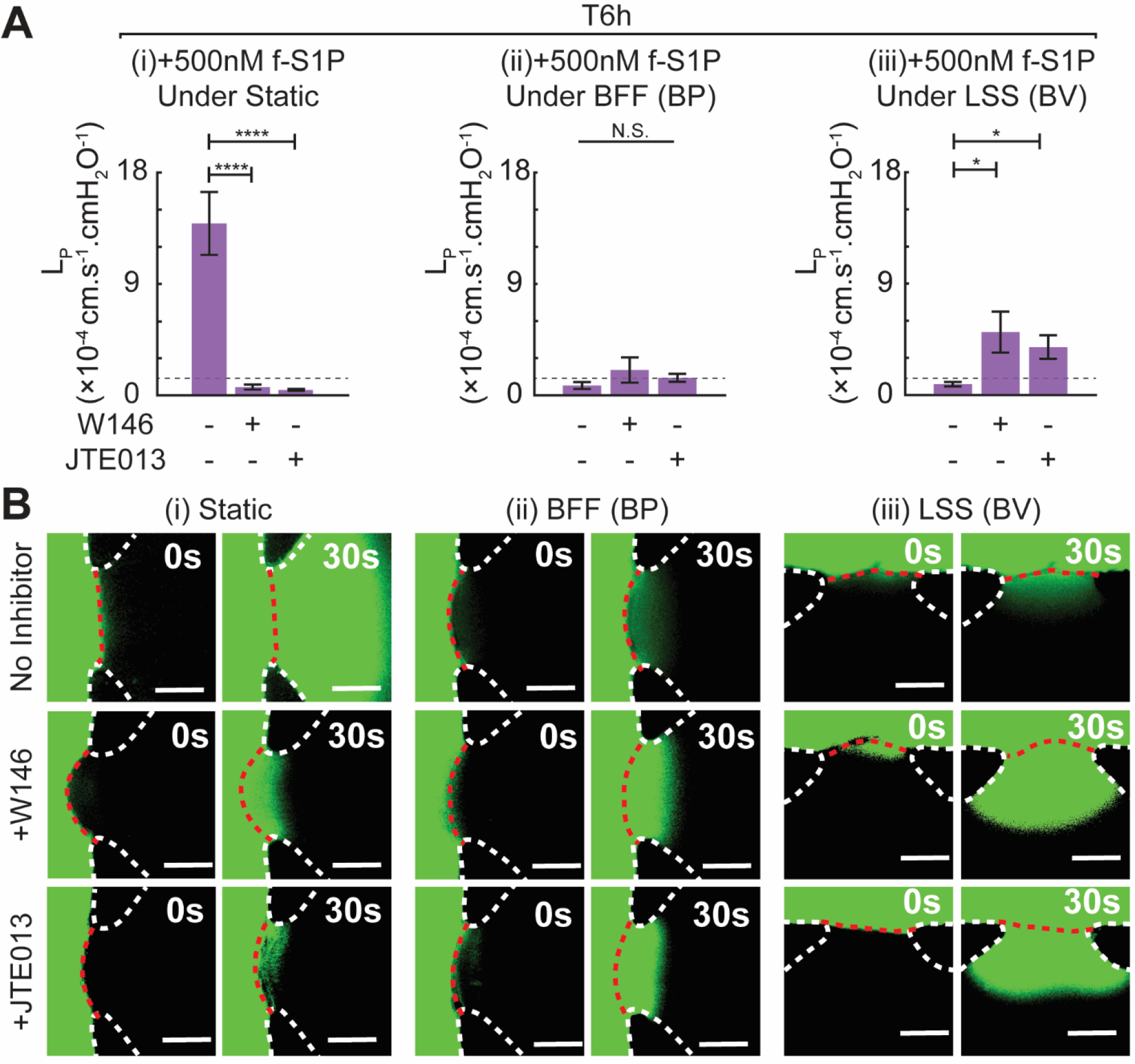
Application of LSS, but not BFF, causes significant increase in *L*_*P*_ when S1PR1 or S1PR2 signaling is inhibited. (A) Quantitative bar-graph plot of HUVEC *L*_*P*_ in response to selective blocking of S1PR1 or S1PR2 signaling followed by treatment with 500nM S1P for 6 hours under (i) static, (ii) when co-applied alongside BFF and (iii) when co-applied alongside LSS. (B) Representative epi-fluorescence images of FITC-Dextran extravasation rate to measure *L*_*P*_ at BP and BV following treatment under each experimental condition. Scale bars are 50µm. *: p < 0.05, ****: p < 0.0001.

In the BP region, pretreatment with W146 or JTE013 followed by co-stimulation with 38 dyn.cm^-2^ BFF and 500nM f-S1P for 6 hours caused a modest but not statistically significant increase in *L*_*P*_ compared to co-application of f-S1P with BFF for 6 hours. W146: 2.04×10^−4^ ± 1.02×10^−4^ cm·s^-1^·cmH_2_O^-1^, *p*>0.05 and JTE013: 1.41×10^−4^ ± 0.32×10^−4^ cm·s^-1^·cmH_2_O^-1^, *p*>0.05 (**Figure 3Aii,Bii**). These results indicate that blocking either S1PR1 or S1PR2 signaling did not significantly affect BFF mediated vessel stabilization. In contrast, in the BV region, pre-treatment with W146 or JTE013 followed by co-application of 3 dyn cm^-2^ LSS with 500nM f-S1P for 6 hours caused a significant increase in *L*_*P*_ compared to co-application of f-S1P with LSS for 6 hours. W146: 5.09×10^−4^ ± 1.65×10^−4^ cm·s^-1^·cmH_2_O^-1^, *p*<0.05 and JTE013: 3.87×10^−4^ ± 0.96×10^−4^ cm·s^-1^·cmH_2_O^-1^, *p*<0.05 (**Figure 3Aiii,Biii**). These results suggest that the enhancement in endothelial barrier function due to LSS is dependent on S1PR1 and S1PR2 signaling.

### 2.4. Association of S1P with albumin carrier transiently promotes vessel stabilization

The effects of S1P on vascular barrier function is known to be carrier dependent,^[12c, 34]^ and one of the primary carriers of plasma S1P is albumin.^[13a]^ Therefore, we measured the effect on HUVEC *L*_*P*_ due to S1P reconstituted with bovine serum albumin (a-S1P) and compared the response with f-S1P under static conditions (**Figure 4**). Treatment with 500 nM a-S1P for 6 hours did not result in a significant difference in *L*_*P*_ (1.69×10^−4^ ± 0.36×10^−4^ cm·s^-1^·cmH_2_O^-1^, *p*>0.05) compared to baseline *L*_*P*_ **(Figure 4A)**. However, the *L*_*P*_ measurement for a-S1P was ∼88% lower than f-S1P at 6 hours of treatment (1.69×10^−4^ versus 13.88×10^−4^ cm·s^-1^·cmH_2_O^-1^, *p*<0.0001; **Figure 4A, B**). These results demonstrate clearly that the effects of S1P on *L*_*P*_ are dependent on association with albumin as a carrier molecule. It is noted that the preparation of f-S1P and a-S1P stock solutions require different buffer conditions (see the Experimental Section). We performed a set of control experiments and confirmed that the differential effects of f-S1P and a-S1P on HUVEC *L*_*P*_ were not due to the different buffer preparations (**Figure S2**).

**Figure 4.**
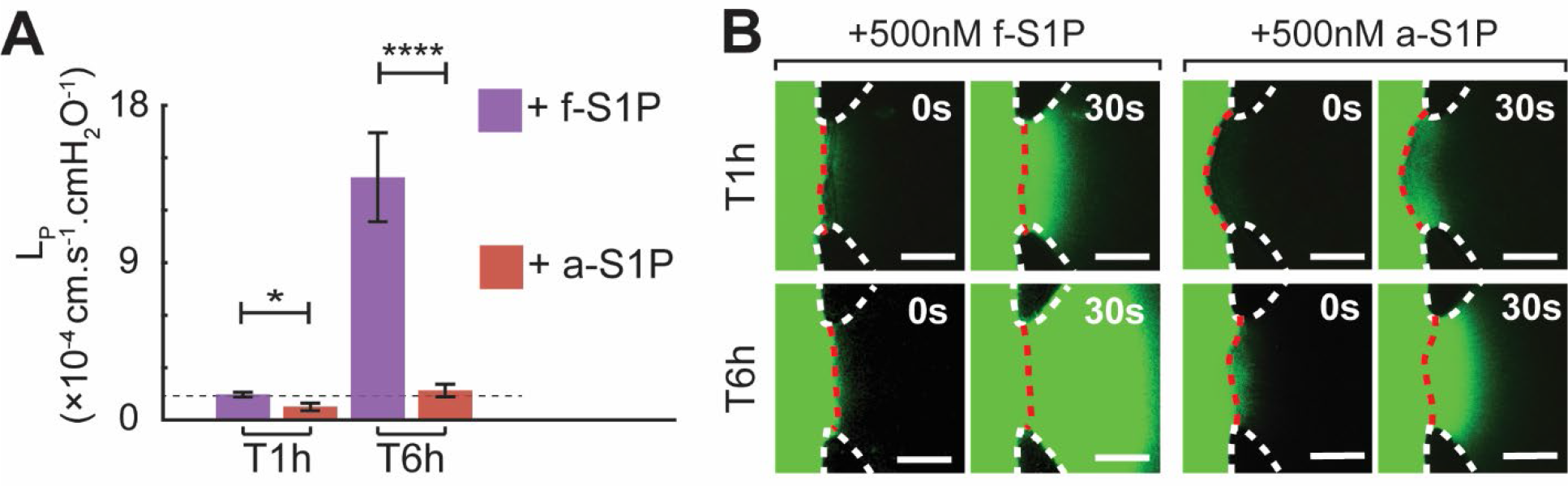
Effect of albumin associated S1P (a-S1P) on endothelial hydraulic conductivity (*L*_*P*_d). (A) Quantitative report on the time-dependent effect of a-S1P on *L*_*P*_ compared to f-S1P. (B) Representative epifluorescence images of the BP aperture depicting the extravasation rate of FITC-Dextran during *L*_*P*_ measurement following treatment with f-S1P versus a-S1P. *: *p* < 0.05, ****: *p* < 0.0001.

We also measured HUVEC *L*_*P*_ due to treatment with a-S1P or f-S1P for 1 hour under static conditions. These studies were motivated by previous reports that demonstrated the effects of S1P on endothelial permeability being time-dependent. For instance, treatment with a-S1P (150-500 nM) under static conditions caused a peak in the transendothelial impedance, which was indicative of enhanced endothelial barrier function, within 30-60 min. This response was followed by a steady decline in barrier function that equilibrated to control levels by 4-5 hours.^[12c, 35]^ Our observed HUVEC *L*_*P*_ measurements at 1 hour and 6 hours of a-S1P treatment under static conditions were in accordance with these previously reported time-dependent measurements for transendothelial electrical resistance. While treatment with 500 nM a-S1P for 1 hour under static conditions caused a significant decrease in *L*_*P*_ (0.69×10^−4^ ± 0.22×10^−4^ cm·s^-1^·cmH_2_O^-1^, *p*<0.05) compared to baseline *L*_*P*_, the same treatment conditions for 6 hours resulted in no difference in *L*_*P*_ compared to baseline *L*_*P*_ **(Figure 4A)**. In contrast to a-S1P, treatment with 500 nM f-S1P for 1 hour under static conditions did not elicit a significant change in *L*_*P*_ (1.46×10^−4^ ± 0.13×10^−4^ cm·s^-1^·cmH_2_O^-1^, *p*>0.05) compared to baseline *L*_*P*_ (**Figure 4A**). Therefore, *L*_*P*_ for a-S1P was ∼48% lower than f-S1P at 1 hour of treatment (0.76 ×10^−4^ versus 1.46×10^−4^ cm·s^-1^·cmH_2_O^-1^ respectively, *p*<0.05; **Figure 4A, B**).

Next, we examined the role of S1PR1 and S1PR2 in mediating the vessel stabilization effects of 1 hour of a-S1P treatment under static conditions. Selective inhibition of S1PR1 signaling with pretreatment of W146 followed by treatment with 500nM a-S1P for 1 hour abrogated the stabilizing effect of a-S1P (1.69×10^−4^ ± 0.32×10^−4^ cm·s^-1^·cmH_2_O^-1^, *p*<0.05) (**Figure S3**). In contrast, blocking S1PR2 signaling with JTE013 followed by treatment with 500 nM a-S1P did not impact the stabilizing effect of a-S1P (0.97×10^−4^ ± 0.17×10^−4^ cm·s^-1^·cmH_2_O^-1^, *p*>0.05) (**Figure S3**). These findings suggest that the transient vessel stabilization induced by a-S1P treatment (**Figure 4A, B**) requires S1PR1 but not S1PR2 signaling.

## 3. Discussion

Laminar shear stress (LSS) due to blood flow has been reported to influence S1P signaling to the endothelium^[23c]^. However, vascular networks are comprised of branching structures, which generate hemodynamic factors that are distinct from LSS. At the base of vessel bifurcations, blood circulation generates bifurcated fluid flow (BFF), which results in stagnation pressure and average zero shear stress at this location. Yet, the role of BFF and LSS in mediating the effects of f-S1P on endothelial permeability remains unstudied in *in vivo* models because of the challenges in controlling hemodynamic factors and biomolecular conditions while deriving quantifiable metrics for vascular barrier function. Here we provided a deeper understanding of how the flow dynamics associated with blood vessel bifurcations fine-tune S1P-mediated vessel permeability. These studies were enabled by the capabilities of our recently reported microfluidic model of vessel bifurcations, which provides quantitative measurements of the vessel permeability coefficient (*L*_*P*_) in response to well-controlled levels of BFF and LSS.^[28a]^

While plasma-borne, carrier-bound S1P is known to be a vasoprotective and atheroprotective factor,^[13b, 14]^ previous studies have shown that carrier-free S1P (f-S1P) promotes compromised barrier function associated with inflammation and atherogenesis.^[18a, 18c]^ Moreover, elevated levels of extravascular-borne f-S1P have been linked to increased destabilization of the tumor vasculature and is a promoter of pulmonary inflammation.^[36]^ As a baseline, we measured vessel permeability due to treatment with f-S1P under static conditions, where we observed a potent induction in HUVEC permeability. This observation was in accordance with previous reports on the role of f-S1P in promoting inflammatory responses in ECs^[18b, 18c]^ and angiogenic sprouting.^[10b]^ When f-S1P is co-applied with BFF and LSS, however, we observed a significant decrease in HUVEC permeability compared to the condition when f-S1P is applied under static conditions. These findings highlighted the prominent role of both BFF and LSS in suppressing the induction of vessel permeability by f-S1P.

We previously reported on the vessel stabilizing effects of BFF and LSS, where we observed significantly decreased *L*_*P*_ for these conditions compared to the static control condition.^[28a]^ Since the magnitudes for BFF (38 dyn cm^-2^), LSS (3 dyn cm^-2^), and duration of treatment (6 hours) in this previous study were the same as the present study, we can make a proper comparison. We observed that the presented values for HUVEC *L*_*P*_ when f-S1P is co-stimulated with BFF and LSS were slightly higher compared to our previously reported HUVEC *L*_*P*_ measurements for the same BFF and LSS conditions but in the absence of f-S1P (∼0.43×10^−4^ cm·s^-1^·cmH_2_O^-1^ and ∼0.35×10^−4^ cm·s^-1^·cmH_2_O^-1^, respectively).^[28a]^ Notably, the observed that changes in *L*_*P*_ in response to treatment with f-S1P under static or perfusion conditions were not attributed to changes in expression of VE-cadherin at the EC junctions. While there is evidence for S1P-induced assembly of VE-cadherin adherent junctions^[15a]^, S1P has been shown to transiently enhance the endothelial barrier integrity through Rho-dependent cell spreading and independent of VE-cadherin binding.^[37]^ Our present findings suggest that f-S1P induced changes in HUVEC L_P_ under static or perfused condition is not dependent on VE-cadherin expression. However, further studies are needed to reveal the mechanism for f-S1P dependent alteration in endothelial junction structure leading to changes in endothelial permeability.

A key area of interest in endothelial mechanosensing is identifying signaling molecules that are also regulated my mechanical forces (i.e. mechanosensors). For instance, we previously reported that the decreases in endothelial permeability by BFF and LSS were both dependent on the nitric oxide (NO) signaling pathway.^[28a]^ However, the mechanisms by which ECs discern between different physical forces (e.g. BFF and LSS) are not known. Thus, a major finding from our study is that vessel stabilization due to LSS, but not BFF, is dependent on S1PR1 or S1PR2. Under perfusion, selectively blocking either S1PR1 or S1PR2 signaling did not cause a significant change in HUVEC *L*_*P*_ in response to co-treatment with f-S1P and BFF. In contrast, under LSS, blocking either S1PR1 or S1PR2 signaling caused a significant increase in *L*_*P*_. Previous *in vivo* observations on vascular hypersprouting in S1PR1 knock-out mice support our observation on increased L_P_ in response to co-stimulation with f-S1P and LSS at BV if S1PR1 signaling is blocked.^[23c, 38]^ However, these *in vivo* observations did not distinguish between regions of high LSS versus the bifurcation points in terms of level of hypersprouting. Moreover, while endothelial S1PR1 has been implicated as mechanosensitive to LSS in promoting vessel stabilization,^[23c]^ our findings suggest a novel mechanosensitive role for S1PR2 in response to LSS in coordinating endothelial barrier function.

Blood flow (i.e. LSS in particular) enhances transcription and protein level of S1PR1,^[23a, 23c]^ and there is evidence of ligand independent activation of S1PR1 that leads to vessel stabilization^[23c]^. However, there are no previous reports on whether BFF elicits ligand independent activation of S1PR1. Therefore, further studies are required to elucidate the mechanism by which LSS and BFF inhibit vessel destabilization by f-S1P. In addition, we note that in contrast to our findings, there are previous reports of combined S1P treatment and LSS inducing angiogenic sprouting.^[39]^ Yet, it is important to consider that these observations were based on higher levels of LSS (greater than ∼6 dyncm^-2^) compared to our study. Thus, our findings demonstrate that the magnitude of applied hemodynamic factors should be considered when evaluating the effects on S1P-dependent vascular permeability.

Previous studies have highlighted the essential role of protein carriers for S1P (e.g. HDL, albumin) in regulating S1PR1-mediated endothelial barrier integrity.^[40]^ These carriers can engage specific endothelial co-receptors, which are not activated when treated with f-S1P.^[40]^ Therefore, we also compared effect of f-S1P on HUVEC *L*_*P*_ to when it is associated with albumin carrier (a-S1P). In contrast with the observed increase in *L*_*P*_ induced by f-S1P, treatment with a-S1P under static condition caused a transient enhancement of the HUVEC barrier that returned to baseline levels after 6 hours. Furthermore, enhancement of HUVEC barrier by a-S1P was mediated by S1PR1 signaling and was independent of S1PR2 signaling. These findings suggest biased activation of S1PR1 over S1PR2 by a-S1P, which were in agreement with previously reported dependence of S1P signaling on its carrier.^[34b]^

In terms of the pathophysiological relevance of our findings, it is well established that flow patterns in tumor associated vasculature are highly abnormal with regions of low flow or flow stasis.^[41]^ For instance, Yuan et al. showed that maximum velocity in tumor-free pial venules are one to three orders of magnitude greater than tumor-associated vessels of comparable diameter.^[42]^ It is worth noting that the estimated Reynolds (Re) number in the tumor-free pial venules was ∼0.3, ^[42]^ which is within the range of the equivalent Re number in the BV region tested in this study. Furthermore, heightened sphingosine kinase activation is present in the cells of the tumor microenvironment, which results in upregulation of stroma-derived S1P production.^[43]^ Therefore, our findings suggest that the absence of ordered and physiological hemodynamic conditions in the tumor-associated vasculature combined with elevated S1P from the tumor stroma may be contributing factors to pathological angiogenesis.

## 4. Conclusion

This study presents to our knowledge the first report on the importance of hemodynamic factors associated with vessel bifurcations (i.e., BFF and LSS) in regulating S1P-mediated vascular barrier function. Moreover, our findings provide a detailed understanding of the role of S1PR1 and S1PR2 in coordinating changes in HUVEC *L*_*P*_ induced by f-S1P. These findings were enabled by the versatility of the described *in vitro* microfluidic model. Future studies using this model will further enhance the understanding of hemodynamic factors in co-regulating S1P signaling, with potential relevance to vascular barrier function and protection against inflammatory, atherogenic, and oncogenic disorders.

## 5. Experimental Section

### Chemical reagents

To prepare stock solution of sphingosine-1-phosphate (S1P), S1P (Cayman) was dissolved in 1X Dulbecco’s phosphate-buffered saline (DPBS) (Corning) with 0.3 M NaOH (Sigma-Aldrich) at 10 mM. To make stock solution of S1P associated with fatty acid-free Bovine Serum Albumin (BSA) (Sigma) carrier proteins, S1P (Avanti Polar Lipids) was resuspended in a methanol: water solution (95:5 volumetric ratio), and heated to 65 °C with sonication to form a 0.5 mgmL^-1^ S1P solution. This solution was then dried with dry nitrogen stream. The dried S1P residue was dissolved in 1X PBS with 4 mgmL^-1^ fatty acid free BSA to a final concentration of 125µM S1P. Stock solution of W146 (Cayman) was prepared by dissolving in DMSO at 50mM. Stock solution of JTE-013 (Cayman) was prepared by dissolving in DMSO at 10mM.

### Microfluidic platform

The microfluidic platform was fabricated and implemented as previously described^[28a]^. Briefly, the microfluidic device consists of a 1300µm wide parent microchannel that symmetrically branches into two downstream daughter microchannels that are 500µm in width. Moreover, the parent microchannel branches around a central extracellular matrix compartment (400µm wide) that encloses the collagenous hydrogel while allowing for direct interaction between the endothelial cells seeded in the microchannels and the collagen matrix at the base of the bifurcation point and within the daughter microchannels. Direct contact between ECs and collagen matrix was enabled by including 100µm wide gaps in the PDMS barrier (referred to as apertures) that separates the ECM compartment from the microchannels. To form individual microfluidic devices, polydimethylsiloxane (PDMS) solution was prepared by mixing silicon elastomer base and curing agent (Ellsworth Adhesives) at a weight ratio of 10:1, and was cast on a silicon master that featured the 50µm in height monolithic microfluidic patterns microfabricated using SU-8 photolithography. The PDMS microdevices were irreversibly bonded on glass slides using plasma treatment and sterilized with UV light prior to casting the collagenous hydrogel.

### Type I collagen hydrogel preparation

The 3-D extracellular matrix (ECM) was comprised of a 3mg/mL collagen type I from rat tail (Corning) solution that was casted and polymerized within the central compartment of the microdevice. A basic solution was prepared using 10X DPBS (Thermo Fisher) and 1 M NaOH (Sigma-Aldrich) to titrate the collagen solution pH to 7.4. To facilitate the adhesion of the endothelial cells and formation of a confluent monolayer on each ECM interface, the collagenous solution was supplemented with human fibronectin (Corning) to a final concentration of 10 µgmL^-1^ fibronectin. The final 3mgmL^-1^ collagenous solution with pH≈7.4 was incubated on ice for ∼10min prior to injection into the ECM microchannel. Following the injection, the cast microdevices were incubated at 37 °C to enable proper polymerization of the collagen fibers.

### Preparation of HUVECs

Human umbilical vein endothelial cells (HUVECs) (Lonza) were cultured using endothelial growth media (EGM) (Lonza) in a cell culture incubator at 37 °C with 5% CO_2_. To seed the microdevices, HUVECs (passage numbers between 5 and 10) were rinsed with 1X DPBS without Mg/Ca (Thermo Fisher), followed by incubation with 0.05% Trypsin-EDTA (Thermo Fisher) for 45 seconds at 37 °C to detach the ECs from the cell culture flask. The detached cells were re-suspended in fresh EGM and prepared for seeding the microdevices. The microdevices that included the polymerized collagen matrix were flushed with 10µgmL^-1^ human fibronectin solution (Corning) diluted in 1X DPBS and incubated for 90 min at 37 °C. Fibronectin-coated microfluidic channels were then flushed with EGM and incubated overnight at 37 °C prior to seeding the HUVECs into the perfusion channels. The HUVECs were removed from the cell culture flask with trypsin and re-suspended in EGM at ∼ 40,000 cellsμL^-1^. The microfluidic channels were then injected with the cell suspension and incubated overnight at 37 °C to facilitate the formation of microdevices fully coated with HUVEC monolayer.

### Pharmacological antagonization of S1P receptors

To pharmacologically antagonize S1P receptor 1, the microdevices seeded with HUVECs were incubated with EGM supplemented with 10µM W146 for 3 hours at 37 °C following the overnight incubation that enables confluent coverage of the device with ECs. The devices were then flushed with EGM prior to the application of each experimental test condition. To pharmacologically antagonize S1P receptor 2, the microdevices seeded with HUVECs were incubated with EGM supplemented with 200nM JTE-013 for 30 min at 37 °C following the overnight incubation that enables confluent coverage of the device with ECs. The devices were then flushed with EGM prior to application of each experimental test condition.

### Immunofluorescence

Microfluidic devices were flushed three times with 1X DPBS and incubated with 4% paraformaldehyde (Sigma-Aldrich) in 1X PBS for 20 min at room temperature following each experimental test condition. The microfluidic devices were then flushed 3 times with 1X DPBS and incubated with blocking buffer, which consists of 5% BSA (Sigma-Aldrich) and 0.1% Triton X-100 (Sigma-Aldrich) for 1 hour. Next, the devices were rinsed 3 times with 1X DPBS, and incubated for 90 min with Alexa647 conjugated anti-human VE-cadherin primary antibody (Life Sciences). Then, the devices were flushed 3 times with 1X DPBS followed by incubation with DAPI (Sigma-Aldrich) diluted in double distilled water by 1:1000 for 3 min to stain for HUVEC nuclei. The devices were finally flushed 3 times with 1X PBS prior to imaging.

### Image acquisition

Before and after treatment under each condition, the HUVECs were imaged using phase contrast imaging. Furthermore, epifluorescence imaging was performed using epifluorescence microscopy (473nm excitation / 488nm emission, TS100, Nikon) with a 10X air objective to monitor the transendothelial transport of FITC conjugated 10 kDa Dextran (Sigma Aldrich). Timelapse epifluorescence imaging was performed at 1s intervals for up to 5 min to capture the dynamic transendothelial transport of the fluorescent tracer. The timelapse epifluorescence images were analyzed using MATLAB to quantify endothelial hydraulic conductivity as previously reported^[28a]^. Confocal microscopy was performed on the stained microdevices using a laser scanning confocal scope (A1R, Nikon) with a 40X oil immersion objective to examine the interendothelial junction structure at the bifurcation point (BP) and in each branched vessel (BV) aperture. A laser type light source was used to excite DAPI (blue) and Alexa 647 conjugated VE-cadherin antibody (far red). VE-cadherin expression at each aperture was examined *en face* by reconstituting a 3-D rendering of the immunofluorescence signal based on multiple confocal images (0.5µm per image slice).

### Statistical analysis

Numerical values reported in the results section represent the mean ± the standard error of the mean of at least three replicates for each experimental test condition. Two-sided student t-tests were used to report the statistical significance between each pair of experimental test condition for *L*_*P*_. Levels of significance were reported using the following: * indicates *p-value* < 0.05, ** indicates *p-value* < 0.01, and *** indicates *p-value* < 0.001.

## Acknowledgements

This work was supported by the National Heart Lung Blood Institute (R01HL141941), American Heart Association (15SDG25480000), and the Center for Emergent Materials, an NSF-MRSEC, grant DMR-1420451, the Center for Exploration of Novel Complex Materials, and the Institute for Materials Research. E.A. and K.K.R acknowledge OSU Presidential Fellowships. G.B.S. acknowledges funding from the Undergraduate Summer Research Program of The American Heart Association Great Rivers Affiliate, a Barry M. Goldwater Scholarship, and an Ohio State University (OSU) Comprehensive Cancer Center Pelotonia Fellowship. M.M.M. acknowledges funding from an Ohio State University (OSU) Comprehensive Cancer Center Pelotonia Fellowship. The authors acknowledge the OSU Campus Microscopy and Imaging Facility (CMIF) for assistance with confocal microscopy. This facility is supported in part by grant P30 CA016058 from the NCI. We thank Peter Beshay, Chia-Wen Chang, and Jonathan Adorno for helpful discussions.

