## Supporting Information for "Endothelial Barrier Function is co-regulated at Vessel Bifurcations by Fluid Forces and Sphingosine-1-Phosphate"

**Number of figures: 3**

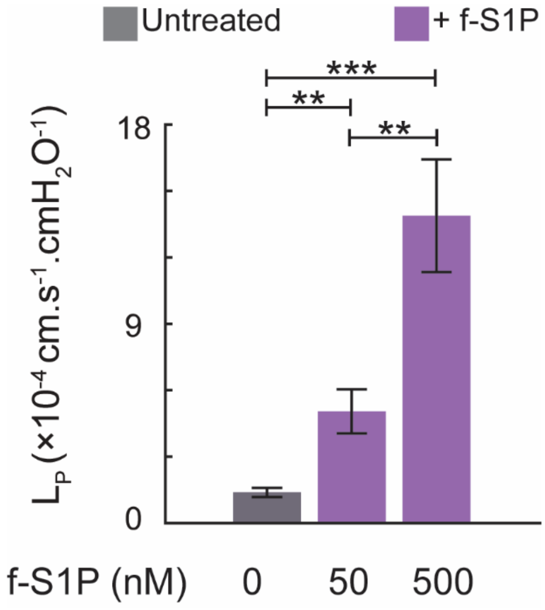


**Figure S1 – Concentration-dependent role of f-S1P in increasing HUVEC permeability under static condition.** Treatment for 6 hours with f-S1P under static condition resulted in a dose-dependent and significant increase in HUVEC $L_P$. **: $p<0.01$, ***: $p<0.001$.

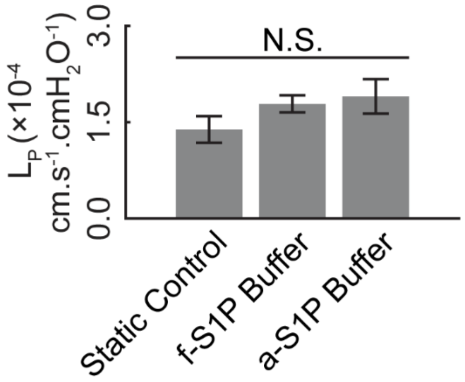


**Figure S2 – Effect of S1P control buffer on endothelial permeability.** Quantitative representation of the endothelial hydraulic conductivity response to the buffer withwhich f-S1P and a-S1P were prepared. To test for the effect of S1P stock buffer solution, we treated the HUVECs for 6 hours with culture media containing f-S1P and a-S1P stock solution buffer diluted to final concentration used in the studies. Treatment with solutions of either buffer did not induce a significant change in $L_P$ compared to static control condition, thereby verifying that the observed changes in $L_P$ mediated by f-S1P and a-S1P were not due to the S1P preparation buffer.

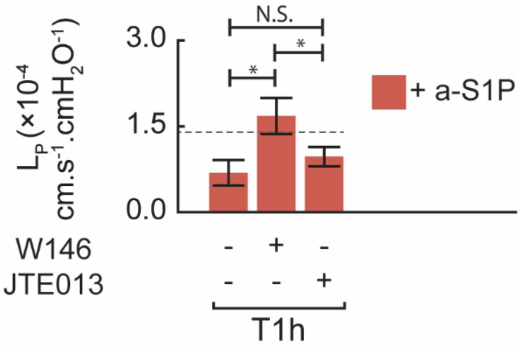


**Figure S3 – Role of S1PR1 and S1PR2 signaling in mediating temporal stabilization of HUVEC monolayer by a-S1P under static condition.** Blocking S1PR1 signaling with W146 significantly inhibited the observed stabilizing effect of treatment with a-S1P for 1 hour. In contrast, blocking S1PR2 signaling with JTE013 did not cause a significant change in L*p* compared to the treated case without any S1P receptor inhibition. These finding suggest a predominant role for S1PR1 signaling, but not S1PR2, in mediating temporal stabilization of the HUVEC monolayer by a-S1P. *: $p < 0.05$.
